# Perturbation Recovery Time Identifies Subtle Human Balance Impairments and Features

**DOI:** 10.1101/2025.06.26.661833

**Authors:** Jiaen Wu, Michael Raitor, Takara Everest Truong, C. Karen Liu, Steven H. Collins

## Abstract

Falls are a leading cause of injury and growing healthcare costs, particularly in aging populations. However, early-stage balance impairments often remain undetected until severe problems arise. Quantitative assessment of human balance therefore remains a critical challenge. Here, we introduce Perturbation Recovery Time, a novel state-space-based balance metric inspired by nonlinear dynamic system theory that quantifies the duration required for gait dynamics to consistently return to a steady-state neighborhood following a perturbation. Unlike conventional approaches based on steady-state walking, this framework evaluates balance through externally induced perturbations that reveal control mechanisms not observable during unperturbed gait. Using healthy participants with experimentally induced impairments, we identified key balance-related features, including the anterior-posterior center-of-mass–center-of-pressure distance, whole-body angular momentum in the frontal and sagittal planes, and vertical center-of-mass position and acceleration. These features capture the core mechanisms of balance control, including foot placement, inverted pendulum dynamics, push-off control, and trunk regulation. Perturbation Recovery Time reliably detected within-subject changes in balance associated with controlled impairments in sensory input, motor coordination, and movement consistency. These findings demonstrate that Perturbation Recovery Time provides a quantitative and sensitive measure of balance, offering a framework for early detection of balance impairments.

## I. Introduction

Falls are a leading cause of injury, disability, and mortality, especially among older adults [1]. Quantitative assessment of human balance is therefore critical for identifying fall risks, informing early interventions, and guiding the design of assistive devices [2]–[5] to enhance walking confidence and reduce falls [6], [7].

Balance assessment has been studied for more than 170 years [8], evolving from functional task-based evaluations to quantitative gait-based metrics. Functional task-based assessments, such as the Short Physical Performance Battery [9] and the Mini Balance Evaluation Systems Test [10], are widely used in older adults [11] and neurological patients [10], [12]. While effective in predicting fall risks, these tests often exhibit floor and ceiling effects, limiting their ability to detect small balance capability differences among highly functional or severely impaired groups. Steady-state metrics, such as step width variability, have been linked to fall risk [13], [14]. Non-linear dynamics-based measures, including Lyapunov exponents and Floquet multipliers, quantify local dynamic stability by linearizing gait dynamics around a limit cycle [15]. Therefore, they primarily capture small fluctuations during steady-state walking and are not designed to characterize balance under perturbations [16]. This limitation is important because real-world falls often arise from unanticipated perturbations, such as trips, slips, or sudden changes in direction [17], [18]. Some balance impairments, such as vestibular dysfunction [19] or muscle weakness [20], [21], may not be evident during steady-state gait. To directly probe balance control, external perturbations have therefore been used to elicit balance responses [20], [22], [23]. While several studies have used time-to-recovery-like measures [24], [25], their ability to detect balance changes has not been demonstrated. Accurately quantifying walking balance remains an open challenge.

Quantifying balance requires biomechanical feature variables that are informative about balance. Different aspects of human movement related to balance have been reported, such as joint angles [26], muscle activations [27], center of mass kinematics [28], [29], center of pressure [28], [30], or angular momentum [31], [32]. However, there is no clear consensus on which biomechanical state variables serve as the sensitive indicators of human balance capability [33].

This study aims to develop a sensitive and reliable metric for quantifying human balance and to identify the most informative balance-related features. We hypothesized that analyzing human responses to external perturbations could reveal key aspects of balance control and enable the development of metrics for real-world balance assessment. In this study, we defined balance as the ability to maintain and restore stable walking following disturbances. To probe this capability, we used controlled perturbations to elicit measurable responses and quantify recovery dynamics. We developed a state-space-based metric, Perturbation Recovery Time, defined as the duration required for gait dynamics to consistently return to a steady-state neighborhood following a perturbation. Although derived from recovery dynamics, this metric assesses overall balance capability rather than recovery alone. Perturbation Recovery Time is computed from state trajectories (Fig. 1 and 2). To establish a steady-state baseline, we model walking as a limit cycle in state space and define a steady-state neighborhood that captures natural variability (Fig. 2A). Together, they provide a reference for detecting gait deviations and determining recovery after perturbations. Since the limit cycle depends on the selected state variables, we systematically evaluated and identified biomechanical features that best characterize balance dynamics and evaluated their contributions to the metric.

**Fig. 1.**
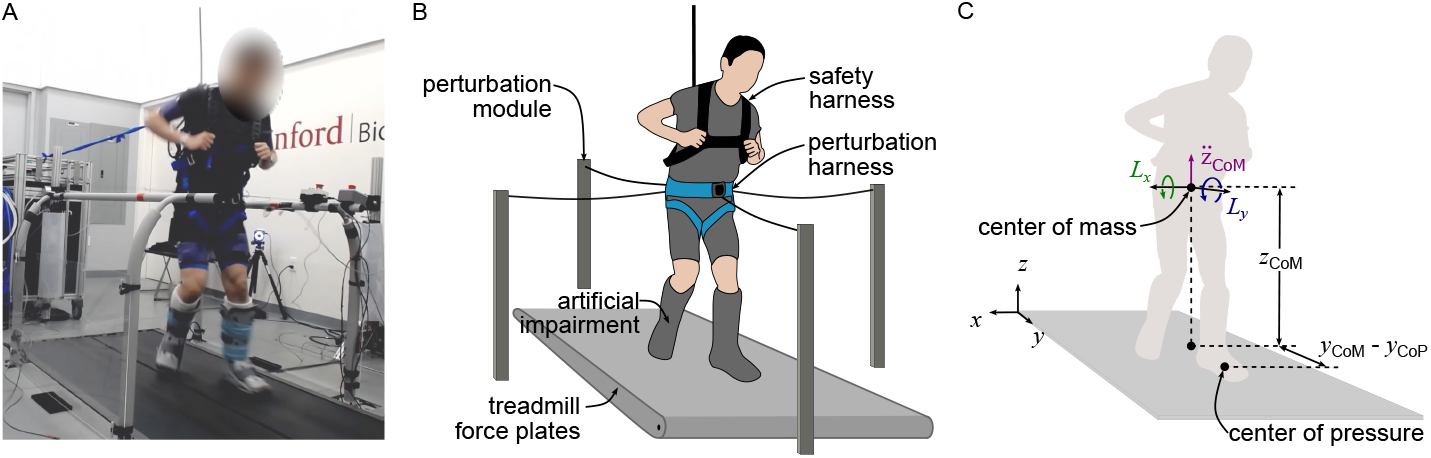
Experimental setup and key balance features identified. (A) An example photograph of a participant with artificial impairment wearing a perturbation harness, a safety harness, and motion capture markers while walking on a treadmill following perturbations. (B) Schematic illustration of the perturbation system setup used to induce controlled perturbations during walking. (C) Schematic representation of the five key features identified as most informative to walking balance capability: the anterior-posterior distance between the center of mass and the center of pressure (*y*_CoM_ − *y*_CoP_), the vertical acceleration of the center-of-mass 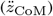, the whole-body angular momentum in the frontal (*L*_*y*_) and sagittal (*L*_*x*_) planes, and the center-of-mass vertical position (*z*_CoM_).

**Fig. 2.**
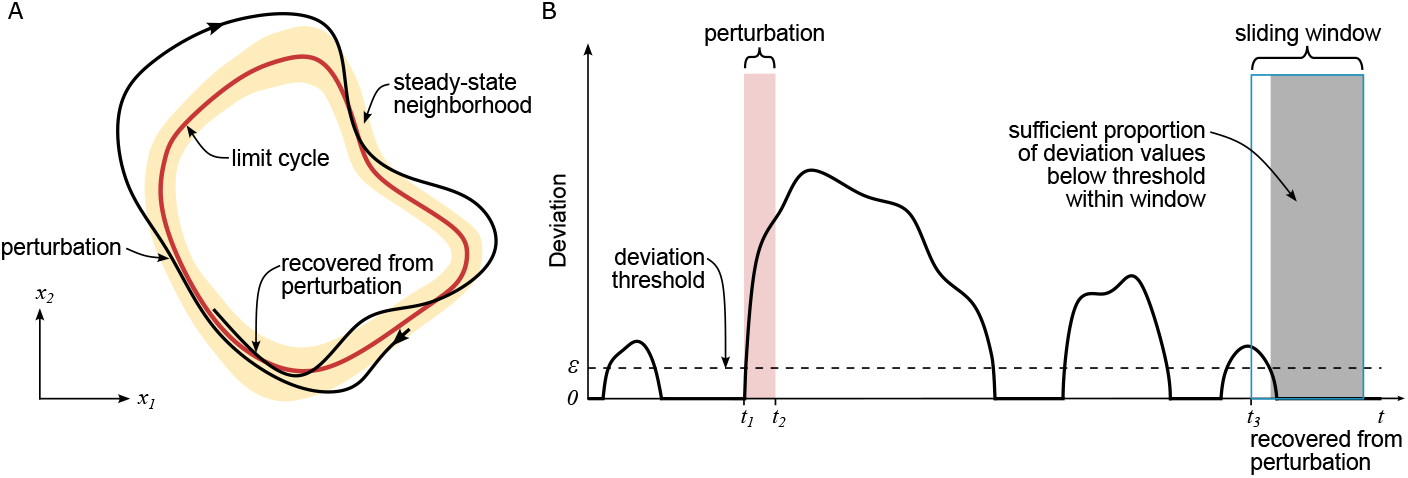
Schematic illustration of the limit cycle and designed deviation metric underlying Perturbation Recovery Time. (A) The limit cycle (red line) and its surrounding steady-state neighborhood (yellow shading) in the state space (*x*_1_, *x*_2_). The black line depicts a walking trajectory that deviates from the limit cycle following a perturbation, with an arrow indicating the direction of travel. (B) The deviation metric over time. The perturbation begins at *t*_1_ and ends at *t*_2_. The deviation threshold *ε* is defined as the 80th percentile of steady-state deviation. Perturbation Recovery Time is identified as the timestamp *t*_3_ when a sufficient proportion of deviation values within a sliding window fall below the threshold.

To develop and validate this approach, we designed a study with healthy individuals whose balance capability was selectively challenged using artificial impairments targeting sensory input, motor control, and movement consistency. These manipulations created controlled and repeatable variations in balance capability while preserving within-subject comparability. We used artificial impairments instead of clinical populations for two reasons. First, they avoid confounding factors commonly present in patients, such as comorbidities, medication effects, and heterogeneous pathology, enabling precise control of impairment type and severity. Second, they provide a well-controlled and low-risk setting to rigorously validate the technical properties of the proposed metric before extending it to clinical populations. We collected motion data during perturbed walking and developed an optimization framework to identify informative balance features and metric parameters. Rather than training a classifier, this framework constructs a physically interpretable metric of recovery dynamics. To ensure robustness and avoid circular validation, the resulting metric was evaluated using subject- and impairment-holdout tests, sensitivity analyses, and label permutation tests.

## II. Methods

### A. Study design

This study involved ten participants (4 females; age = 24-31 years; mass = 68 ±13 kg; height = 1.71 ±0.09 m), each completed a total of 64 walking trials at a speed of 1.25 m/s, involving controlled perturbation administered across four walking conditions, four perturbation directions, two perturbation magnitudes, and two repetitions per unique combination.

To create controlled and repeatable differences in balance capability, the trials included a control condition (normal shoes) and three artificial impairments, each designed to probe a specific sensorimotor subsystem relevant to human balance. Ankle braces simulated impaired motor coordination by mechanically limiting the ankle motion and increasing joint stiffness. It reduced the speed, precision, and efficiency of ankle strategies used for balance control. This strategy has been used to observe motor deficits in conditions such as ankle osteoarthritis or age-related declines in distal neuromuscular control [23]. Vision occlusion (eyes blocked) removed visual feedback, thereby degrading the spatial awareness of body position and movement, and increasing reliance on proprioceptive and vestibular inputs. This approach has been used to simulate sensory inaccuracy observed in older adults and people with sensory impairments [34], [35]. Pneumatic jets introduced motor noise and execution errors by delivering air bursts randomly starting at 0%, 25%, 50%, or 75% of the swing phase, with durations varying from 25% to 100% of the swing phase. This intervention increased variability in foot placement and decreased movement precision. It simulated coordination deficits such as tremors or neuromuscular noise observed in Parkinson’s disease or cerebellar ataxia [36].

Perturbations were applied using the Bump’em system [37], which generated forces at the pelvic level, approximately aligned with the body’s center of mass to minimize induced rotational effects (Fig. 1A-B). Each perturbation was initiated at the mid-stance phase of the left foot and lasted for 300 ms. Two perturbation magnitudes were used, corresponding to 7.5% and 15% of body weight (equivalent to impulse integrals of 2.25% and 4.5% body weight seconds, respectively). Perturbations were applied in four directions (anterior, posterior, left, and right). Each unique combination of walking condition, perturbation direction, and magnitude was repeated twice. The perturbation order was fully randomized across trials and participants to minimize potential learning, anticipation, and systematic order effects. Full-body kinematics were recorded at 100 Hz using an 11-camera optical motion capture system (Vicon Motion Systems, Yarnton, UK) with a Plug-in Gait marker set with additional markers on the upper arms, thighs, and shanks. Ground reaction forces (Bertec, Columbus, OH, USA) and perturbation forces were measured at 1000 Hz. Surface electromyography was collected at 1000 Hz from eight muscles per leg (gluteus maximus, gluteus medius, rectus femoris, vastus lateralis, semitendinosus, medial gastrocnemius, soleus, and tibialis anterior) using Delsys sensors. Additional experimental details are provided in [16].

This protocol was approved by the Stanford Institutional Review Board (IRB-57846).

### B. Data processing

Gait cycles were segmented with heel strike events. Center-of-mass kinematics and whole-body angular momentum were computed in OpenSim [38], using musculoskeletal models scaled to each participant’s anthropometry. Marker locations were adjusted to match the ankle brace configuration. Inverse kinematics was performed using markers unaffected by these interventions to ensure consistent pose estimation. The added mass of the ankle braces was incorporated via scaling with preserved mass distribution, maintaining consistent segment inertial properties without changing geometry.

To minimize the influence of station-keeping during treadmill walking, we removed low-frequency drift in the center-of-mass position (*<* 0.3 Hz) associated with slow positional adjustments. This was achieved by estimating the drift using a fourth-order low-pass Butterworth filter (0.3 Hz cutoff) and subtracting it from the original signal, followed by low-pass filtering at 6 Hz to retain gait dynamics. Note that the station-keeping did not affect the state variables ultimately used in the Perturbation Recovery Time metric. The center of mass kinematics, whole-body angular momentum, and center of pressure data were low-pass filtered using a fourth-order Butterworth filter with a selected cutoff frequency of 6-15 Hz, depending on the state variable. Finally, state variables for each trial were normalized by their means and standard deviations obtained from the last five steady-state strides of that trial, ensuring comparability between different features.

### C. Perturbation recovery time definition

We defined Perturbation Recovery Time as the minimum duration required for a person’s gait to consistently return to the neighborhood of its steady-state limit cycle following a perturbation. The calculation involved three main steps (Fig 2): (1) characterizing steady-state gait as a limit cycle and its steady-state neighborhood in the state space (Fig. 2A), quantifying the deviation from this steady state after a perturbation (Fig. 2B), and (3) determining the time when this deviation consistently falls below a threshold (Fig. 2B).

Once balance features were identified (Section II-E), they were concatenated into a multidimensional state vector representing gait state at each time instant. This defines a state space that provides a geometric framework for characterizing steady-state gait, perturbation-induced deviations, and recovery.

#### 1) Steady-state gait characterizatio

Steady-state gait was modeled as a limit cycle in a state space defined by selected biomechanical balance features (Section II-E). The limit cycle, denoted as 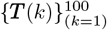, represented the average trajectory of the state vector over one gait cycle, reflecting the inherent consistency in locomotion. It was sampled at 100 points, which was sufficient to capture the nuances of the gait cycle while maintaining computational efficiency. Here, ***T*** (*k*) denoted the state vector on the limit cycle at stamp *k* ∈ {1, 2, …, 100}. The natural stride-to-stride variability was captured by the standard deviation trajectory of multiple steady-state walking cycles, denoted as 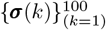. To compute this information, we collected at least 80 steady-state strides from each perturbation direction and magnitude for each experimental condition and participant. The steady-state neighborhood 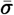, defined as the average standard deviation trajectory, represented the inherent fluctuation during unperturbed walking.

#### 2) Deviation metric design

To quantify transient recovery after perturbations, we designed a deviation trajectory metric ***D***(*t, k*^*⋆*^) that applied weighted penalties to selected state variables’ deviations that exceed the predefined walking consistency. After a perturbation, balance states ***S***(*t*) deviated from the limit cycle. To quantify this, we first identified the sample index *k*^*⋆*^ of the closest state on the limit cycle to the current state ***S***(*t*) using a weighted distance metric:

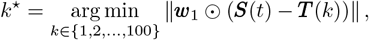

where ***w***_1_ was a weight vector defining the relative importance of each state variable, ⊙ denoted the element-wise multiplication, ‖· ‖ denoted the 2-norm of the vector, and *t* denoted time. Bold letters represent vectors, and nonbold letters represent scalars. Importantly, *k*^*\**^ does not represent a temporal or phase-aligned index in the gait cycle, but the point on the limit cycle that is closest to the current perturbed state in the constructed state space. This geometric projection allows recovery to be assessed without assuming preserved gait timing, which may be disrupted immediately following a perturbation. While event-based or phase-aligned matching may be appropriate for steady-state analyses, it relies on reliable gait event detection and preserved temporal structure, assumptions that may not hold during transient perturbation recovery. The deviation trajectory metric ***D***(*t, k*^*⋆*^) was then calculated as the magnitude of the significant deviations beyond the steady-state neighborhood:

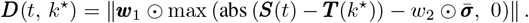

Here, abs (***S***(*t*) *−****T*** (*k*^*⋆*^)) computed the element-wise absolute difference between the current state and the closest state on the limit cycle. This difference was compared to the corresponding steady-state neighborhood 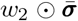, where the scalar *w*_2_ denoted a neighborhood scaling factor that uniformly defined the tolerance range for all features based on their natural variability 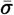. The element-wise maximum function max(·, 0) ensured only deviations exceeding this tolerant neighborhood range were incorporated into ***D***(*t, k*^*⋆*^). Finally, these significant deviations were weighted by the feature importance vector ***w***_1_ and aggregated into a single scalar value ***D***(*t, k*^*⋆*^). This formulation independently controls the relative importance (***w***_1_) of biomechanical state variables and the tolerance to natural gait variability (*w*_2_), resulting in a physiologically consistent and numerically robust metric. The separation of ***w***_1_ and *w*_2_ is more general and has a better interpretation than using one single parameter, because a feature’s importance in defining balance is not necessarily proportional to its natural variability (e.g., some features might be noisier). Moreover, no overfitting of this design was observed during extensive validation tests (Section III-B). During steady-state walking, values of ***D***(*t, k*^*⋆*^) remained consistently low across all participants, confirming it captures primarily perturbation-induced deviations rather than natural gait variability.

#### 3) Perturbation Recovery Time determination

Perturbation Recovery Time was determined as the time duration from the start of perturbation to the earliest time point *t*^*⋆*^ when the deviation ***D***(*t, k*^*⋆*^) consistently remained low, indicating a return to steady-state gait. This was formalized using a sliding window of duration *w*_*3*_:

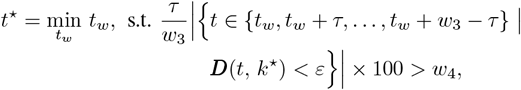

where *w*_3_ ∈ {*τ*, 2*τ*, 3*τ*, …} denoted the sliding window size, *τ* denoted sampling time, *w*_4_ ∈ [0, 100] represented the window proportion threshold, meaning that at least *w*_4_% states within this window size remained within the neighborhood of the limit cycle. |·| denoted the cardinality counting the number of samples within the sliding window {*t*_*w*_, *t*_*w*_ + *τ*, …, *t*_*w*_ + *w*_3_− *τ*} where ***D***(*t, k*^*⋆*^) was below the deviation threshold *ε. ε* was a predefined threshold that reflected the deviation of steady-state walking. We set this threshold *ε* to the 80th percentile of steady-state ***D***(*t, k*^*⋆*^) values, calculated from last five strides in a trial after recovery. This value was selected after testing percentiles from 70th to 95th for its optimal tradeoff between sensitivity to recovery and robustness to natural gait variability. The Perturbation Recovery Time *t*^*⋆*^ was the minimum time *t*_*w*_ (in units of seconds) that satisfies the given condition.

### D. Optimization framework

To identify informative biomechanical features and hyperparameters for the Perturbation Recovery Time metric, we formulated a constrained optimization problem. The goal was to determine a parameter set 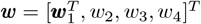 that defines the state-space representation and deviation metric used to compute the Perturbation Recovery Time. The objective function promotes consistent separation between conditions across perturbation trials, while preserving robustness to natural gait variability. This optimization operates at the level of metric construction (i.e., feature weighting and thresholding) rather than enforcing outcome differences.

The cost function was designed to minimize the summation of two terms subject to ∥***w***_**1**_∥ _1_ = 1. The first term (weighted by *C*_1_ *>* 0) promoted a longer average Perturbation Recovery Time for each impaired condition *e* ∈ {AB, EB, PJ} com-pared to the normal condition (*e* = NS) across perturbation directions. Here, the indicator function 𝟙 (·) counted successful comparisons. It returned 1 if the inside statement was true and 0 otherwise. The second term rewarded differences in the Perturbation Recovery Time between normal and impaired conditions. The exponential function exp(·), with a weight parameter *C*_2_ *>* 0, grows rapidly with increasing differences. It was designed to reward consistent to moderate differences while preventing outliers from dominating the results. The *l*_1_ normalization constraint on ***w***_**1**_ ensured the feature weights sum up to one. *t*_*e,s,p*_ was the Perturbation Recovery Time for experimental condition index^1^ *e* ∈ {NS, AB, EB, PJ}, subject index *s* ∈ {1, 2, …, 10}, and perturbation index *p* ∈ anterior, left, right, posterior. NS, AB, EB, and PJ represented normal shoes, ankle braces, eyes-blocked, and pneumatic jets, respectively. *n*_*e*_ was the number of experimental conditions, *n*_*s*_ was the number of subjects, and *n*_*p*_ was the number of perturbation directions. This formulation resulted in a multi-level optimization problem, which was non-convex, discontinuous, and high-dimensional, arising from the indicator function, sliding window evaluation, and multi-objective trade-offs. We solved it using the Covariance Matrix Adaptation Evolution Strategy (CMA-ES), a stochastic optimizer well suited for such problems [39].

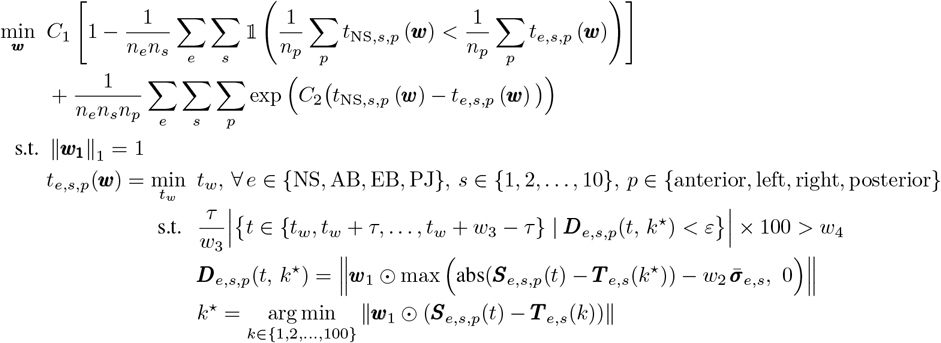

### E. Balance feature identification

We used an iterative selection and validation process to identify the most informative features for assessing human balance capability. This involved evaluating candidate features for defining limit cycles and assessing how accurately the resulting Perturbation Recovery Time metric distinguished between normal and impaired conditions across participants.

We began with a candidate list of kinematic and kinetic variables previously used in the literature as related to fundamental balance mechanisms and control [26]–[32]. This feature set captured whole-body dynamics. It included three-dimensional position, velocity, and acceleration of the whole-body center of mass and pelvis; three-dimensional whole-body angular momentum and central inertia; the horizontal displacement and velocity difference between the center of pressure and the center of mass; three-dimensional trunk kinematics including position, velocity, acceleration, angular velocity, angular acceleration, and trunk’s displacement and angle relative to the center of mass. We then systematically evaluated all possible feature combinations, testing subsets of 1 to 8 features at a time. For each combination, we optimized feature weights and assessed the Perturbation Recovery Time’s accuracy in distinguishing different balance capabilities. The features that consistently demonstrated the best performance across validation schemes were identified as the most sensitive and reliable for encoding balance mechanisms. These features were subsequently used to define the limit cycle and compute the Perturbation Recovery Time. Note that the role of optimization was not to invent new features, but to identify which of these pre-selected variables were most sensitive for detecting subtle balance changes.

### F. Generalizability and robustness assessment

We conducted a series of tests to evaluate the robustness and generalizability of Perturbation Recovery Time across new subjects, impairment types, and experimental conditions.

#### 1) Parameter sensitivity analysis

To assess the robustness of the optimized parameters 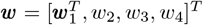, we performed a global sensitivity analysis. We defined a sensitivity range of ±30% around each optimized parameter value (e.g., for a parameter value of 0.20, the tested range was 0.14 to 0.26). For each parameter, we fixed all other parameters at their optimized values and systematically swept the parameter of interest over a uniformly sampled ±30% range with 1% increments. At each point, we recalculated the Perturbation Recovery Time for all trials and evaluated its performance using two metrics: overall accuracy, i.e., the proportion of cases where the mean Perturbation Recovery Time (across all subjects) for the normal condition (NS) was shorter than for each impaired condition (AB, EB, PJ) (n = 3 comparisons per parameter point); subject-specific accuracy, i.e., the proportion of instances where, for each subject, the mean Perturbation Recovery Time across perturbations was shorter in the NS condition than in each impaired condition (n = 10 subjects × 3 impairments = 30 comparisons per parameter point). This one-at-a-time approach allowed us to isolate the individual effect of each parameter on the Perturbation Recovery Time metric’s discriminatory power.

#### 2) Generalizability to new subjects

We assessed the generalizability of the Perturbation Recovery Time metric using both one and two subjects holdout cross-validation, where the metric was trained on a subset of subjects and tested on unseen ones. In the leave-one-subject-out cross-validation scheme, parameters (***w***) were optimized using data from 9 subjects and tested on the held-out subject (10 iterations). We further extended this to a more stringent test (leave-two-subject-out cross-validation) by holding out 45 possible pairs of two subjects, optimizing parameters on the remaining 8 subjects, and testing on the two held-out subjects. The metric’s training and testing accuracies were analyzed for each evaluation.

#### 3) Generalizability to new impairments

To evaluate the generalizability of the Perturbation Recovery Time metric to unseen types of balance impairment, we performed leave-one-impairment-out validation. For each of the three impaired conditions, we held out all corresponding data. Parameters were optimized using data from the normal condition and the two remaining impaired conditions. The optimized metric was then tested on the unseen, held-out impairment.

#### 4) Assessment of perturbation magnitudes

The Perturbation Recovery Time metric was optimized and tested separately for two perturbation magnitudes: large impulses (4.5% body weight-seconds) and small impulses (2.25% body weight-seconds). For each magnitude, the metric’s generalizability was assessed using both one and two subjects holdout cross-validation.

#### 5) Assessment of perturbation directions

We evaluated the robustness of the Perturbation Recovery Time metric to perturbation directions using two-subject holdout cross-validation across multiple datasets: (1) all four directions (anterior, posterior, left, right); (2) combined anterior-posterior directions; (3) combined medial-lateral directions; and (4) individual direction separately. To decouple the effects of directional diversity from data volume, we also trained the model using data from all four directions but with only a single repetition.

#### 6) Performance with feature reduction

To evaluate each feature’s contribution, we systematically removed features from the original set of five and assessed the metric’s performance across all possible feature combinations using two-subject holdout cross-validation. Accuracy was averaged across all cross-validation iterations of all possible feature combinations.

#### 7) Overfitting analysis via label permutation tests

To assess whether Perturbation Recovery Time’s performance was driven by meaningful balance information or model overfitting, we performed two label permutation tests, including one label shuffling test and three label swapping tests, and assessed their performance using two-subject holdout cross-validation. In the label shuffling test, we randomly reassigned condition labels to each trial to destroy any real association between balance conditions and labels, then retrained and tested the metric under these conditions. Performance at chance level (50%) would indicate no overfitting. In the label swapping test, we exchanged labels in pairs between normal and each impaired condition, then retrained and tested the metric’s performance. A substantial performance drop would indicate that Perturbation Recovery Time relied on meaningful balance information rather than superficial condition classifications.

### G. Statistical analysis

Statistical analyses were performed to compare the Perturbation Recovery Time across conditions. Data normality was assessed with the Shapiro–Wilk test and variance homogeneity with Levene’s test. When both assumptions were satisfied, paired t-tests were applied; otherwise, Wilcoxon signed-rank tests were used. A significance level of 0.05 was used. Analyses were performed in Python 3.9 (Numpy 1.21.5, Pandas 1.5.3, Scikit-learn 1.2.2, and MATLAB R2021b.

## III. Results

### A. Identified balance features and optimized parameters

Five features were identified as most informative to balance capability (Fig. 1C): the anterior-posterior distance between the center of mass and the center of pressure (*y*_CoM_*−y*_CoP_), the center-of-mass acceleration in the vertical direction 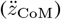, which is proportional to the vertical ground reaction force, the whole-body angular momentum in the frontal (*L*_*y*_) and sagittal (*L*_*x*_) planes, and the center-of-mass vertical position (*z*_CoM_). All five variables contributed substantially to post-perturbation deviations (Fig. 5). The anterior-posterior distance between the center of mass and the center of pressure contributed most across all conditions. The vertical position of the center of mass had the smallest contribution. The neighborhood of the limit cycle was defined using a neighborhood scaling factor of 0.839 applied to the average gait variability. The Perturbation Recovery Time was determined using a sliding window of 2.14 seconds and a window proportion threshold of 79.8%, requiring 79.8% of the deviation signals to remain below the 80th percentile of steady-state walking deviation. This meant that the recovery criteria required a similar steady-state deviation level to be maintained.

### B. Perturbation Recovery Time Differentiates Balance

Perturbation Recovery Time effectively detected changes in balance capability for all artificial impairment types and subjects. At the group level, the normal shoe condition showed the shortest Perturbation Recovery Time (Fig. 3A), indicating the best balance capability, with significant differences compared to each impaired condition (*p* < 0.05). At the individual level, the Perturbation Recovery Time consistently distinguished balance capabilities with longer values under impaired conditions compared to the normal condition (Fig. 3B).

**Fig. 3.**
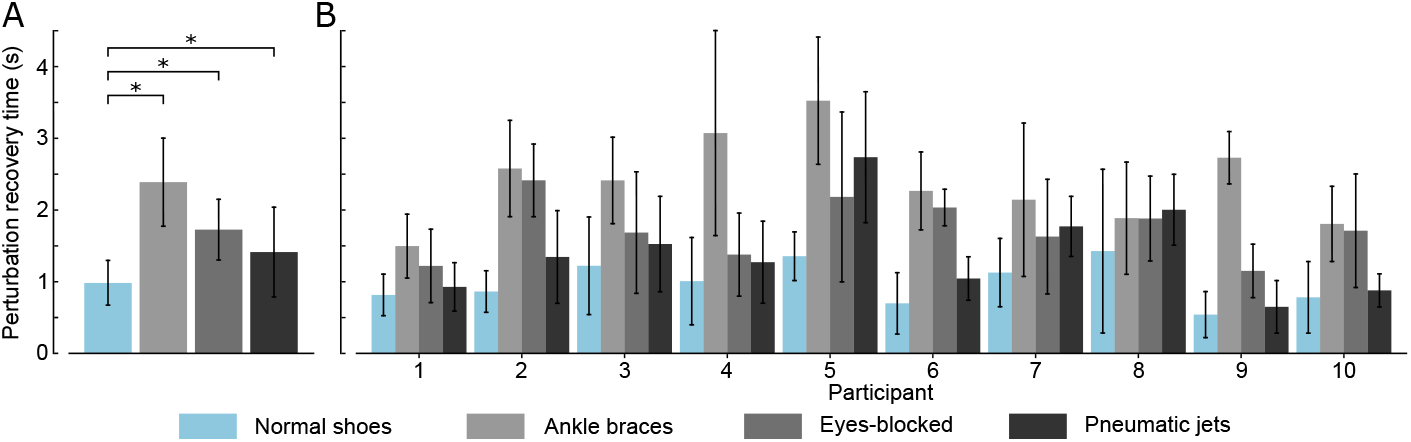
The Perturbation Recovery Time across balance conditions and subjects. (A) Average Perturbation Recovery Time across different balance conditions (n=10). (B) Average Perturbation Recovery Time for each subject (n=8). Error bars represent standard deviations. ^*\**^*p* < 0.05.

#### 1) Sensitivity analysis of the optimization problem

The optimized solution showed low sensitivity to local hyperparameter variations, indicating robustness of the Perturbation Recovery Time metric. Overall accuracy remained near 100% across the entire ±30% variation range for all five feature weights and hyperparameters, including neighborhood scaling factor, sliding window size, and window proportion threshold (Fig. 6). This indicates that the Perturbation Recovery Time metric’s ability to distinguish group-level differences between normal and impaired balance is resilient to changes in hyperparameters. Subject-specific accuracy showed only minor fluctuations, remaining at 100% for parameter variations within ±10%. As parameter variations approached ±30% of the optimized values, performance slightly declined by around 5%. Notably, the window proportion threshold exhibited a critical limit. When it exceeded 93%, accuracy dropped to chance level. This occurs because an overly stringent threshold requires near-perfect consistency within the window, which is rarely met even during steady-state walking. Window percentage thresholds above 100% were infeasible.

#### 2) Performance with feature reduction

The performance of Perturbation Recovery Time declined with decreasing number of features (Fig. 4A). This indicates that each identified feature contributed substantially to capturing the intrinsic balance mechanism, reinforcing the need for a comprehensive feature set to maintain both accuracy and computational efficiency.

**Fig. 4.**
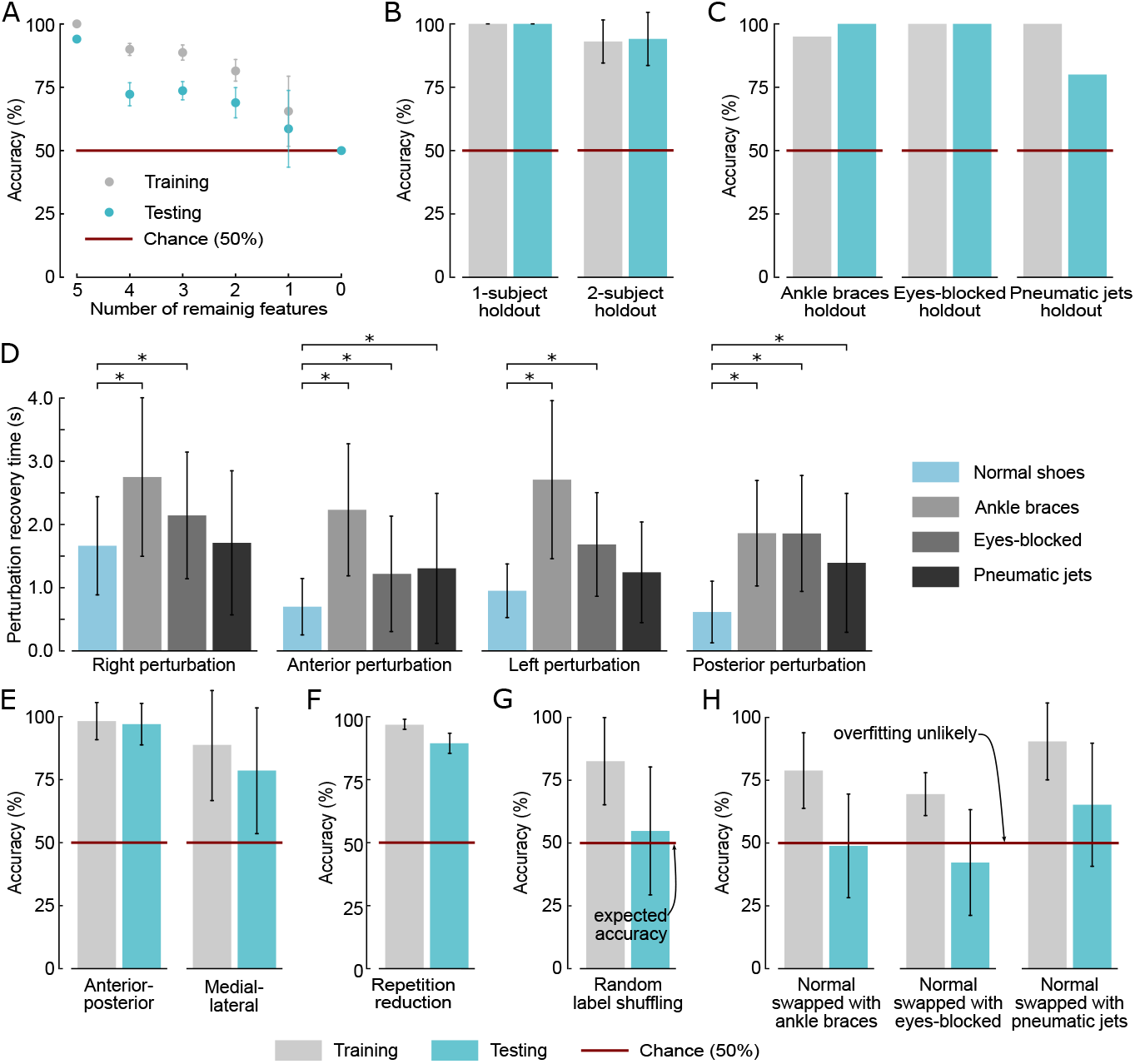
Evaluation of the Perturbation Recovery Time metric’s performance and generalizability. (A) Accuracy of Perturbation Recovery Time as a function of the number of included features using 2-subject holdout cross-validation. (B) Generalizability accuracy when applying the metric to new subjects, evaluated with 1-subject (n=10) and 2-subject (n=45) holdout cross-validation. (C) Generalizability accuracy when introducing new impairment conditions. (D) Perturbation Recovery Time values in four perturbation directions illustrating directional differences in balance recovery (n=20, ^*\**^*p* < 0.05). (E) Accuracy of the Perturbation Recovery Time metric for anterior-posterior and medial-lateral perturbations (n=45). (F) Accuracy of the Perturbation Recovery Time metric when using data from a single repetition across four perturbation directions (n=45). (G-H) Label permutation test for overfitting analysis using 2-subject holdout cross-validation (n=45): (G) random label shuffling test and (H) label swapping test, where labels of the controlled condition (normal shoes) were swapped with one impaired condition. Higher accuracy above 50% indicates potential overfitting, while accuracy values close to or at 50% are preferable. Error bars represent standard deviations.

**Fig. 5.**
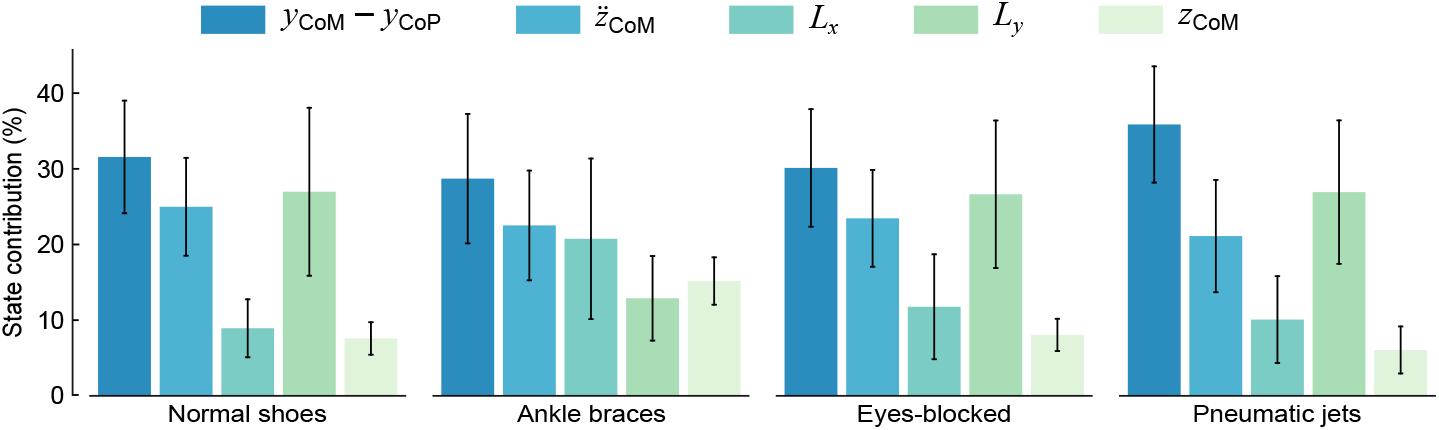
Percentage contribution of state variables to perturbation deviation metric across conditions. Error bars represent standard deviations.

**Fig. 6.**
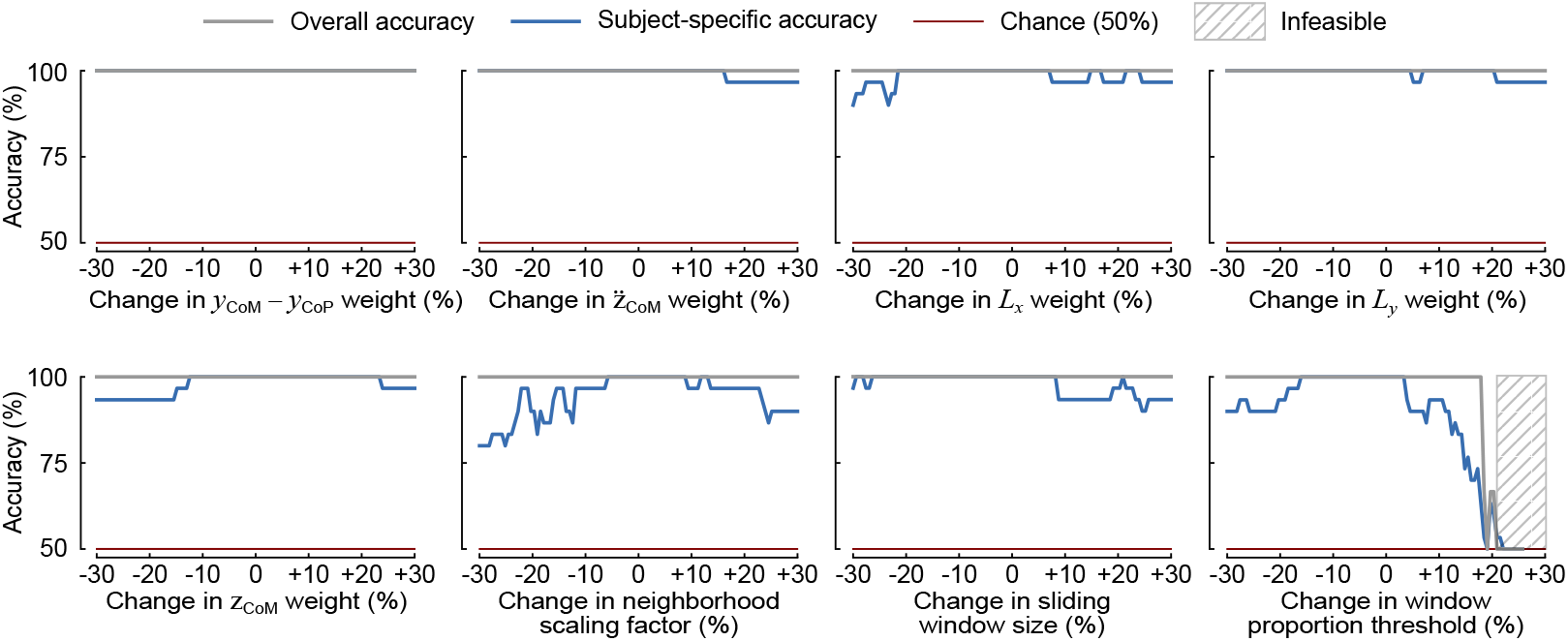
Sensitivity analysis of the Perturbation Recovery Time metric. The impact of varying each optimized parameter individually by ±30% on overall and subject-specific accuracy. Window proportion threshold above 100% is infeasible.

#### 3) Generalizability to new subjects

Perturbation Recovery Time generalized well for new subjects, as evidenced by subject holdout cross-validation experiments (Fig. 4B). It exhibited consistently high performance with an average accuracy of 100% in leave-one-subject-out cross-validation. In the more stringent leave-two-subjects-out validation, training and testing accuracies were 93% and 94%, respectively.

#### 4) Generalizability to new impairment

Perturbation Recovery Time effectively generalized to unseen impairment types. When applied to the held-out conditions, testing accuracy was 100% for both ankle braces and eyes-blocked condition, and 80% for pneumatic jets, yielding a mean testing accuracy of 93% across all three impairments(Fig. 4C).

#### 5) Impact of perturbation magnitudes

Perturbation Recovery Time detected balance changes more effectively with larger perturbations. For larger magnitudes, accuracy reached 100% in single-subject and 94% in two-subject holdout assessments (Fig. 4B). With smaller perturbations, the generalizability declined to 93% (single-subject holdout) and 86% accuracy (two-subject holdout) (Fig. 7A).

**Fig. 7.**
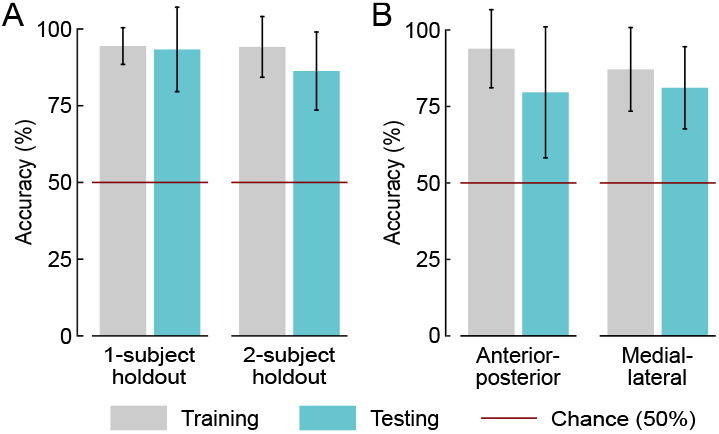
Evaluation of the Perturbation Recovery Time metric. (A) Metric accuracy using 1-subject (n=10) and 2-subject (n=45) holdout cross-validation. (B) Metric accuracy for data from anterior-posterior and medial-lateral directions, respectively (n=45).

#### 6) Impact of perturbation directions

Perturbation Recovery Time effectively differentiated balance impairments across all perturbation directions (Fig. 4D), but performance was direction-dependent. Under larger magnitudes, Perturbation Recovery Time showed higher accuracy for anterior-posterior perturbations (97%) than for medial-lateral perturbations (79%) (Fig. 4E). This directional asymmetry diminished under smaller perturbations, where accuracies were comparable (anterior–posterior: 80%, medial–lateral: 81%; Fig. 7B).

Directional diversity compensated for limited repetitions for optimizing state-variable weightings and hyperparameters used to compute Perturbation Recovery Time. A metric optimized using perturbations from four directions with only one repetition per condition achieved 90% accuracy (Fig. 4F), outperforming metrics optimized using two repetitions from only two directions (anterior-posterior or medial-lateral, Fig. 4E) and approaching the performance of the fully optimized metric using two repetitions across all four directions (Fig. 4B). When optimizing the metric using data from any single perturbation direction, limited data led to overfitting.

#### 7) Overfitting analysis

The Perturbation Recovery Time metric’s ability to differentiate balance capabilities was not due to overfitting. When condition labels were randomly shuffled, the metric’s performance dropped to chance levels (Fig. 4G). This indicated that it had no predictive power without real label associations. Similarly, when pairwise exchanging labels between normal and each impaired condition, the performance declined substantially (Fig. 4H), confirming that Perturbation Recovery Time detects balance changes based on intrinsic balance characteristics rather than classifying conditions.

## IV. Discussion

Perturbation Recovery Time effectively detected small changes in balance capability for each subject. While the artificial impairments did increase gait variability in healthy participants, the magnitude of this variability remained much lower than that typically observed in clinical populations with balance disorders, as quantitatively demonstrated in our previous study [16]. This indicates that the induced deficits were relatively subtle, which demonstrated Perturbation Recovery Time’s ability to identify subtle deficits that conventional assessments may overlook. The values of Perturbation Recovery Time varied by impairment (Fig. 3A): normal shoes (0.98 ±0.29s), eyes-blocked condition (1.73 ±0.41s), ankle braces (2.39 ±0.61s), and pneumatic jets (1.41± 0.62s). These values align with theories on the rapid reflexive and voluntary responses of the human balance system [24], [29].

### A. Influence of perturbation directions and repetitions

Our findings suggest that reliable balance assessments require both diverse perturbation directions and sufficient repetitions. The Perturbation Recovery Time metric performed better with anterior-posterior perturbations than with medial-lateral perturbations when data were limited (Fig. 4E-F). This difference is likely because anterior-posterior responses were more consistent across trials. In contrast, medial-lateral perturbations produced greater response variability, which increased optimization complexity and data requirements. This increased variability may also reflect greater challenge of mediolateral balance control and could provide additional sensitivity to balance deficits and fall risk.

Although we initially expected multiple perturbation directions to be necessary to fully evaluate balance, our results suggest that focusing on a minimal set, such as anterior-posterior perturbations (Fig. 4E), might still provide effective assessments with a large and diverse dataset. Data scarcity remains a challenge when using a single perturbation direction, suggesting the need for future research on minimal yet effective combinations of perturbation directions.

Reducing repetitions while maintaining four perturbation directions preserved the Perturbation Recovery Time metric’s generalizability (Fig. 4F), indicating that directional diversity could partially compensate for fewer repetitions. However, the slight performance drop compared to multi-repetition scenarios emphasizes the importance of repeated trials and sufficient perturbation data for reliable optimization, capturing subtle balance variations, and reducing noise through averaging, thereby improving the metric’s precision and reliability.

### B. Influence of perturbation magnitudes

Larger perturbations improved the differentiation of balance capabilities, while smaller perturbations often produced small deviations that fell within the normal fluctuations of steady-state gait (Fig. 4B and Fig. 7A). Selecting an appropriate perturbation magnitude is therefore critical. Excessive magnitudes may elicit exaggerated responses that do not reflect typical balance mechanisms, whereas small magnitudes may insufficiently challenge balance. Within a certain range, increasing perturbation magnitude may not produce proportional increases in Perturbation Recovery Time. Similar to the step response of dynamic systems’ local behavior, where rise and settling times reflect intrinsic system properties independent of input magnitude [40], Perturbation Recovery Time could reach a plateau where it is determined by the body’s inherent balance mechanisms rather than external perturbation magnitudes. Future studies should systematically characterize this relationship between perturbation magnitude and Perturbation Recovery Time to identify optimal ranges for balance assessment.

The ideal perturbation magnitude may vary across individuals due to their differences in body size, strength, neuromuscular control, and experience with balance tasks. For applications focused on within-subject tracking (e.g., rehabilitation), a personalized magnitude perturbation protocol may be beneficial. Future studies could explore personalized perturbation profiles, such as using a graded approach where perturbation magnitudes are gradually increased until observing a predefined behavioral response, such as the need to take a corrective step, a measurable deviation in trunk kinematics, or the patient’s own perception of instability. However, for cross-subject impairment classification, a standardized magnitude must be used to ensure fair comparisons. Future research could explore normalization methods for perturbation design that enable equitable and sensitive balance assessment across individuals and diverse populations.

### C. Impact of impairments

Three artificial impairments provided different challenges. Among them, pneumatic jets showed the Perturbation Recovery Time closest to the normal condition. This might be because the randomized timing and duration of air bursts during swing phase resulted in more variable and smaller disruptions, with some trials having minimal impact on balance recovery. This also explains why the Perturbation Recovery Time metric’s accuracy dropped to 80% when pneumatic jets were excluded during leave-impairment-out validation. Their under-representation in training data likely prevents the optimization process from fully exploring the parameter landscape, thereby identifying less precise solutions.

The eyes-blocked condition primarily disrupts station keeping on the treadmill, which is a separate control task from perturbation balance recovery. Without visual feedback, participants often wandered on the treadmill, showing difficulties in maintaining position. After filtering out this low-frequency positional drift, participants showed effective balance recovery. This was likely due to their intact alternative sensory systems (e.g., proprioceptive and vestibular) and the predictable walking environment. These observations indicate that vision occlusion represents a subtle impairment for this population. In the ankle brace holdout experiment, test accuracy slightly exceeded training accuracy. While this may seem counterintuitive, it is a known phenomenon in machine learning when training data are noisier or more variable (e.g., pneumatic jets, eyes-blocked), whereas the test data contain clearer, more consistent patterns (e.g., ankle braces, as also reflected in their longer Perturbation Recovery Time values (Fig. 3)) [41], [42]. This result does not indicate overfitting. Instead, it demonstrates the model’s capacity to generalize effectively to clearer impairment patterns unseen during training.

These findings indicate that incorporating diverse and subtle impairments provides richer variability in balance responses, thereby improving optimization of the metric and enhancing its sensitivity and generalizability. This principle could extend to gait analysis [43] and rehabilitation programs.

### D. Perturbation Recovery Time vs. steady-state metrics

The Perturbation Recovery Time metric consistently outperformed commonly used balance metrics, including center of mass displacement, Lyapunov exponents, step width/time variability, foot placement predictability, and lateral margin of stability (Fig. 8), when evaluated on the same perturbation dataset using leave-one-subject-out cross-validation [16]. This performance gap may arise from fundamental differences in what these metrics capture. Conventional metrics typically quantify variability during steady-state walking, which may not reveal balance deficits that do not appear under unchallenged conditions. In contrast, Perturbation Recovery Time directly quantifies the temporal evolution of recovery following destabilizing perturbations, providing a functionally relevant measure of balance control that captures the coordinated neuromuscular response to perturbations. These results indicate that the improved performance of the Perturbation Recovery Time metric is not solely due to the use of perturbations, but also to its time-based, process-oriented formulation that is rooted in dynamical systems theory and synthesizes multiple biomechanically meaningful features.

**Fig. 8.**
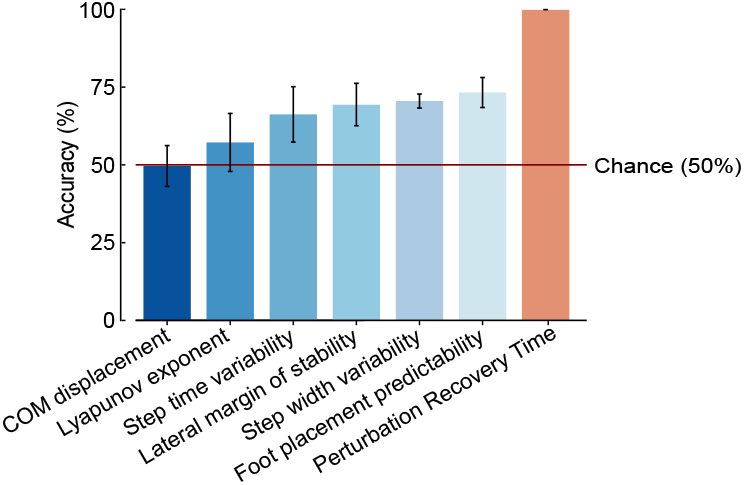
Comparison of balance metric performance on perturbation data evaluated using the one-subject hold-out cross-validation (n=10). Error bars indicate standard deviations across subject holdout evaluations.

While perturbation protocols require greater experimental complexity than steady-state assessment, the increased sensitivity of Perturbation Recovery Time can be valuable for applications requiring high precision. In screening or low-resource settings, steady-state metrics may be sufficient. While for applications where detecting subtle, early-stage decline is important, such as early detection of neuromuscular decline, rehabilitation monitoring, or assistive device tuning, the enhanced sensitivity of Perturbation Recovery Time justifies the additional instrumentation.

### E. Perturbation Recovery Time vs. magnitude-based metrics

We also evaluated several magnitude-based metrics, including various modifications of center-of-mass displacement, but found their discrimination accuracies close to chance (50%) when distinguishing between normal and impaired conditions [16]. As a result, these metrics were excluded from further optimization. The superior performance of the time-based Perturbation Recovery Time metric over magnitude-based measures could be attributed to several reasons. First, Perturbation Recovery Time directly measures the time required to consistently return to steady-state gait, capturing the entire recovery process, including detecting perturbations, initiating corrective actions, and re-establishing steady-state gait. This comprehensive assessment reflects the efficiency of sensory perception, neuromotor control, and muscular capacity, whereas magnitude-based metrics often isolate specific aspects of balance, potentially missing critical phases of balance strategies. Second, Perturbation Recovery Time could be more sensitive to small delays in neural reflexes and muscular responses that indicate subtle balance impairments. Magnitude-based metrics lack a temporal component, making them less effective in detecting these variations. Additionally, Perturbation Recovery Time tracks a continuous recovery process rather than relying on discrete moments, such as center-of-mass excursion [16], [44], which can vary significantly between trials and require more measurements for accurate estimation. Unlike magnitude-based metrics that fluctuate with perturbation intensity, Perturbation Recovery Time could provide a more consistent measure across varying perturbation magnitudes. These findings suggest that a time-based, process-oriented metric like Perturbation Recovery Time could be more informative for assessing balance and fall risks than static measures. This principle could also be applicable to other domains where dynamic system recovery needs to be assessed, such as evaluating stability in robotics or prosthetics.

### F. Informative features for assessing balance

We identified five balance features that effectively characterize balance dynamics, whose deviations following perturbations are used to compute Perturbation Recovery Time. Each feature captures key aspects of gait and balance control, and together they provide a comprehensive description of the mechanisms underlying stable locomotion and recovery.

The anterior-posterior distance between the center of mass and center of pressure reflects foot placement relative to the body’s center of mass. It captures gait mechanics through two main mechanisms: the inverted pendulum model, which maintains a proper distance between the center of mass and center of pressure during steady-state walking, and foot placement control [45], [46], which is an active strategy that adjusts the center of pressure to maintain balance, especially after large perturbations. The vertical position of the center of mass reflects the dynamics of inverted pendulum motion, capturing the energy conversion as the center of mass rises and falls during walking. The vertical acceleration of the center of mass reflects dynamic interactions between the body and the ground, particularly during push-off. Effective push-off generates forward forces to propel the center of mass forward, which is energy-efficient for gait [47] and allows modulation for balance [48]. Whole-body angular momentum reflects torso strategies that facilitate rapid adjustments and corrections of upper-body movements in response to perturbations [22], [31].

We also tested alternative state definitions, such as medial-lateral center-of-mass displacement or foot displacement, but found them less accurate, likely due to their redundancy with other measures, such as whole-body angular momentum in the frontal plane. Additionally, these balance features tended to exhibit greater variability in response to perturbations, suggesting that more data would be needed to consistently extract meaningful balance information. Note that while other features may contain balance-relevant information, the selected set emerged as particularly robust across subjects and impairment types, reflecting their fundamental roles in balance control.

Removing any of these five identified features significantly reduced the performance of the Perturbation Recovery Time metric, confirming the unique contributions of each feature to walking balance. Human gait and balance control are multidimensional processes that depend on the integration of several biomechanical and neurological systems. The interplay of foot placement, inverted pendulum gait characteristics, controlled push-off, and trunk dynamics is crucial for maintaining gait efficiency and balance. The collective integration of these balance features improves the sensitivity of the Perturbation Recovery Time metric in assessing balance, confirming that effective balance assessments require monitoring multiple kinetic and kinematic state variables to capture the multidimensional nature of the control process.

### G. Applications of Perturbation Recovery Time

Perturbation Recovery Time provides a sensitive quantitative measure for detecting within-subject changes in balance capability, with broad potential for clinical and research applications. Clinically, its ability to detect subtle balance impairments enables earlier interventions, therapy evaluation, and individual rehabilitation tracking. In research, the Perturbation Recovery Time metric can guide the design and personalization of assistive devices, such as exoskeletons and prostheses, by allowing objective comparisons of control strategies. It can also be integrated into professional or athletic training to enhance balance performance and sensorimotor adaptability through repeated perturbation-based practice, supporting more effective and individualized training programs.

The Perturbation Recovery Time metric quantifies balance capability regardless of the gait phase in which perturbations occur. While this study utilized stance-phase perturbations, the framework also applies to other perturbation types like swing-phase disturbances (e.g., stumbling) or stance-phase slips. These perturbations engage shared whole-body balance mechanisms captured by the selected balance features. However, the relative contribution of individual features and optimal hyperparameters may vary across perturbation types. Perturbation-specific tuning may therefore be required to preserve metric sensitivity across diverse balance challenges.

### H. Limitations

This study focused on specific perturbation directions and magnitudes in healthy young adults with artificial impairments, which do not replicate all physiological deficits observed in clinical populations. Neurological and age-related balance disorders often involve delayed sensorimotor processing that was not explicitly induced in this study. Such delays would be expected to manifest as prolonged recovery dynamics, which are directly captured by Perturbation Recovery Time. However, when extending this framework to clinical cohorts, population-specific optimization of feature weightings and hyperparameters may be required to account for these differences. In addition, lateral attachment of perturbation cables constrained natural arm swing across all conditions, which may have influenced balance responses despite being consistent across conditions. Future studies should validate the metric in clinical populations through longitudinal studies, while also exploring a wider range of perturbation profiles, developing implementations based on wearable or markerless sensing, establishing population-normative references to enable cross-subject comparisons, and exploring data-driven approaches, such as generative models, to compare distributions of perturbed and unperturbed motion.

## Data and Materials Availability

All data supporting this study are available in the main text or the supplementary materials. The experimental data can be accessed via Dryad (https://doi.org/10.5061/dryad.cnp5hqch3) or through our laboratory server at http://gofile.me/4oVL6/uFylcak0t. The data and code used to produce the results presented in this manuscript are available on GitHub at https://github.com/JiaenWu/PerturbationRecoveryTime.git.

## Acknowledgment

The authors acknowledge Shengling Shi for his thoughtful discussion, feedback, review, and editing of the manuscript.

## Footnotes

1 As ***D, T, S, σ***, 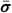 vary across subjects, experimental conditions, and ***D, S*** also depend on the perturbation directions, they are formally written with the subscript indices, e.g., ***T***_*e,s*_ and ***D***_*e,s,p*_. The subscripts are omitted for brevity when no confusion arises.

